## Supplementary figures and images for "The *Mycobacterium tuberculosis* complex pangenome is small and shaped by sub-lineage-specific regions of difference"

### Supplementary Figure S1

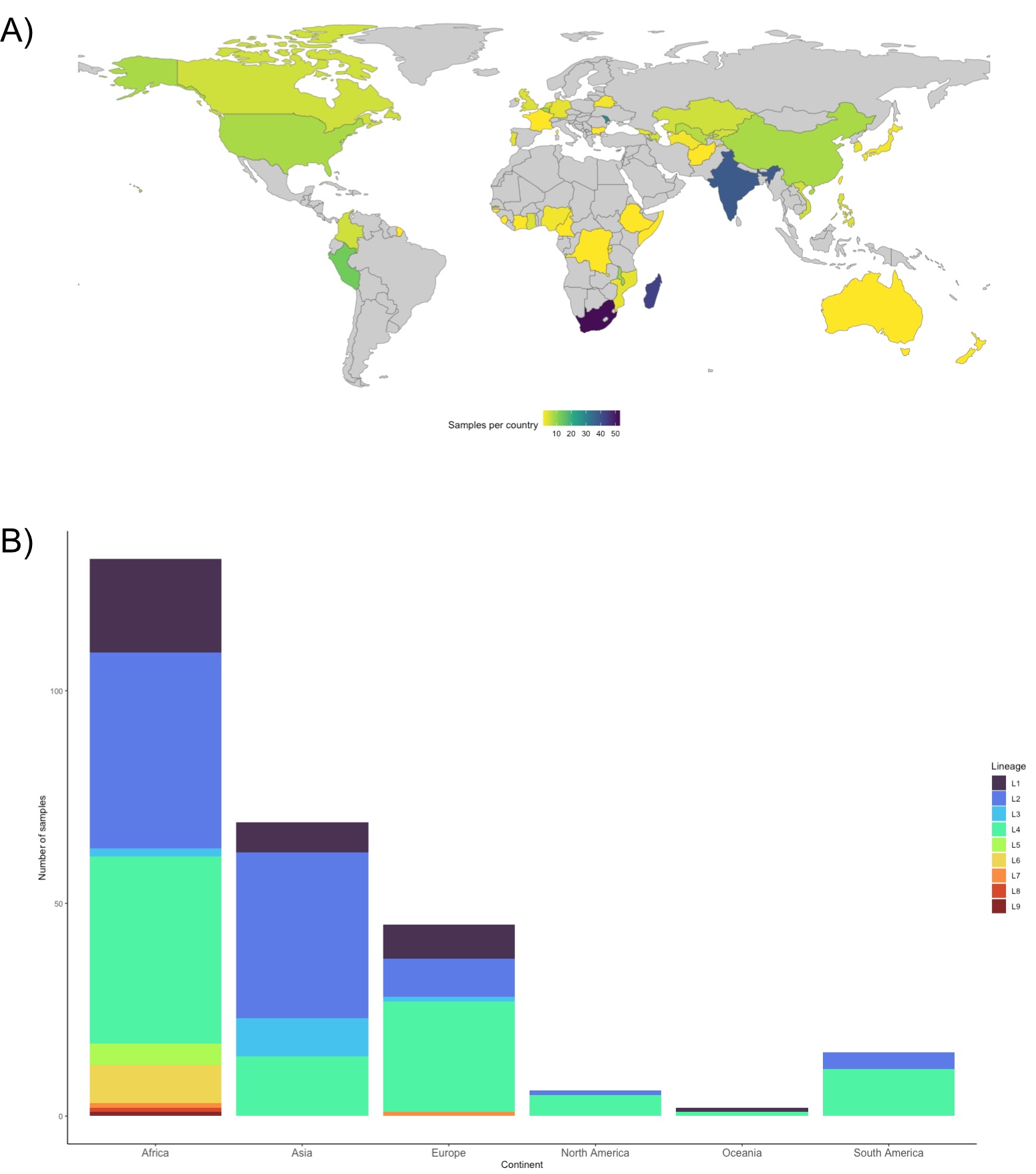

### Supplementary Figure S3

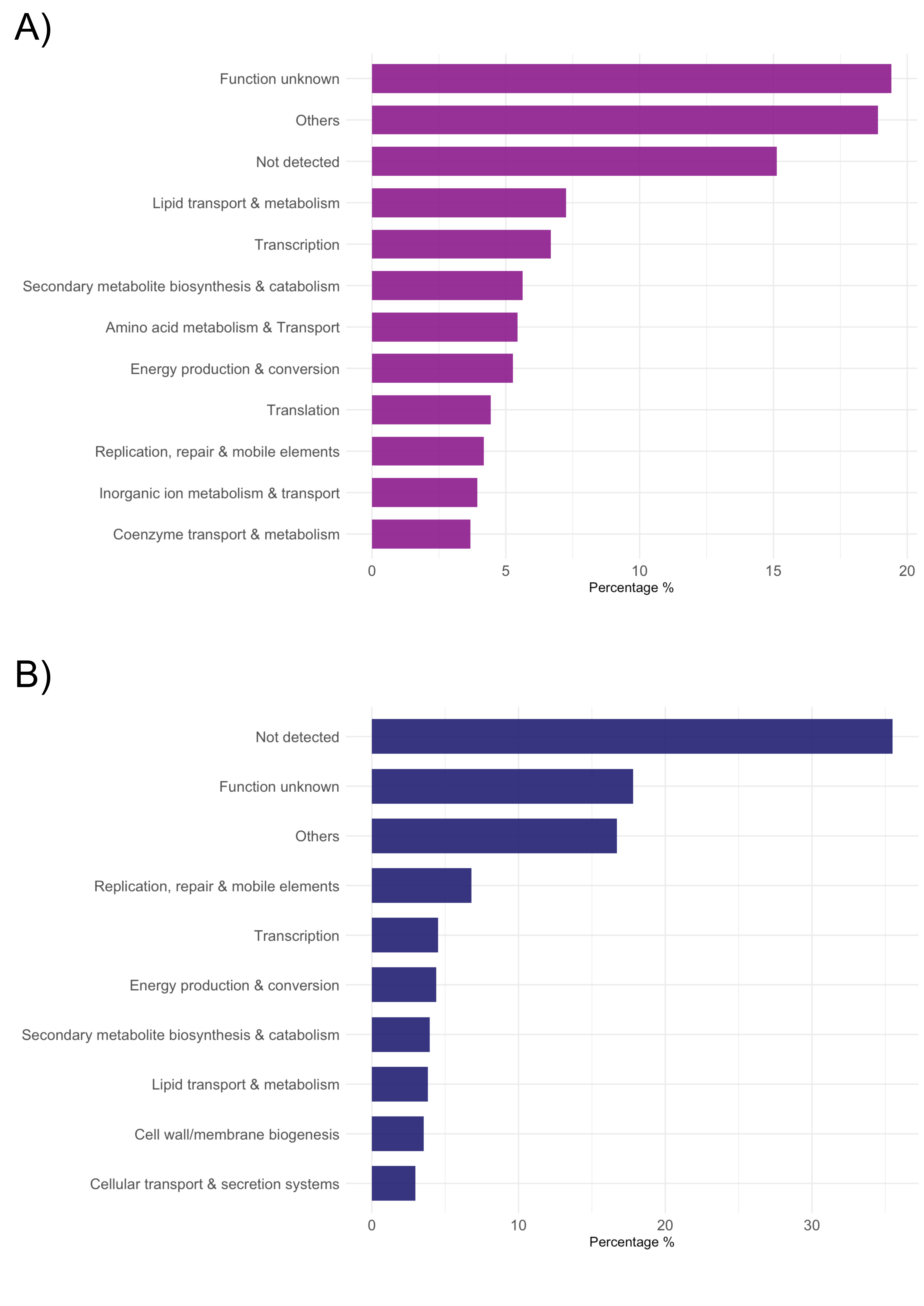

### Supplementary Figure S4

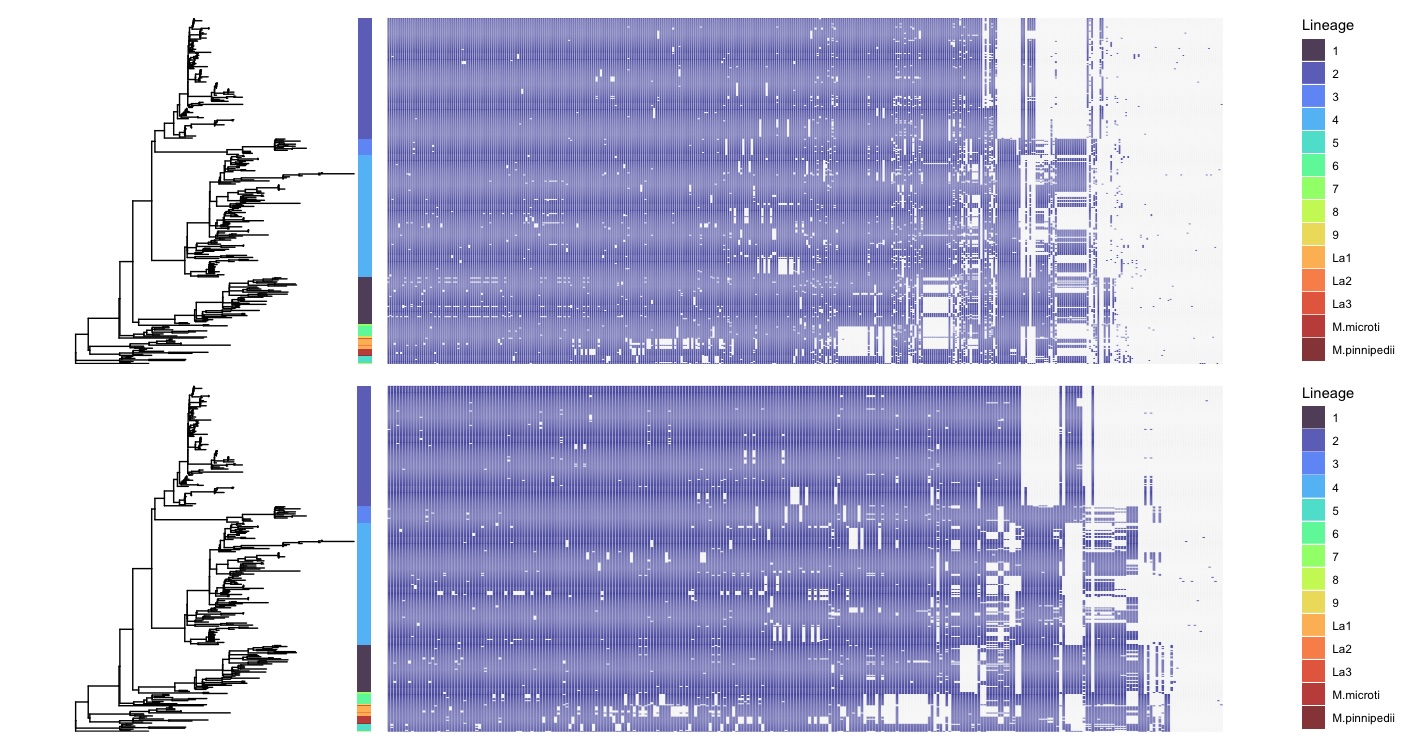

### Supplementary Figure S5

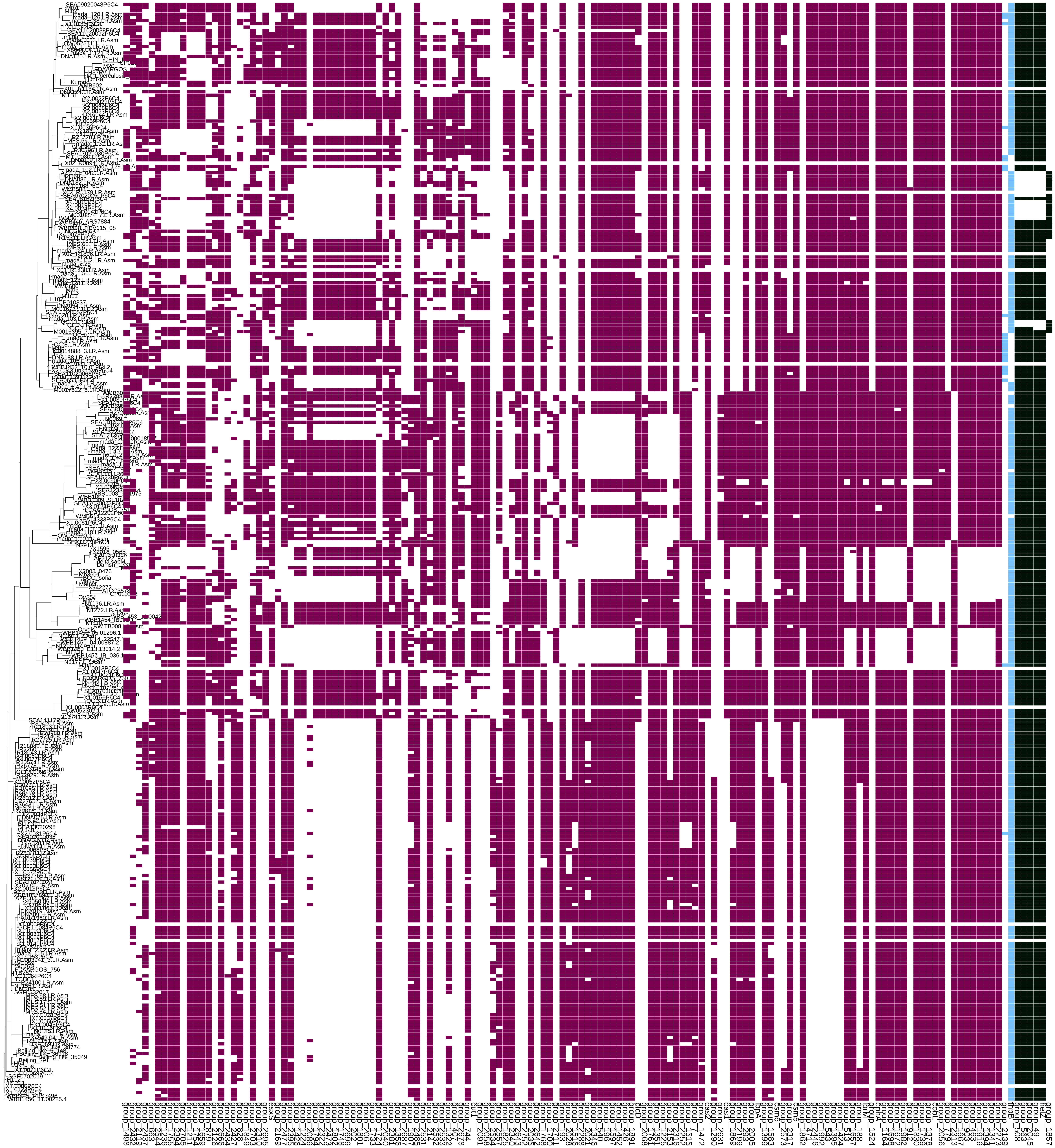
