## Supplementary Figure S2 for "The *Mycobacterium tuberculosis* complex pangenome is small and shaped by sub-lineage-specific regions of difference"

**A) Merged paralogs**:

| **Fit to Heap's law:** | **Intercept** | **alpha** |
|---|---|---|
| | 117.6233 | 2 |

| **Genome Fluidity:** | **Mean** | **Std** |
|---|---|---|
| | 0.008487286 | 0.003941261 |

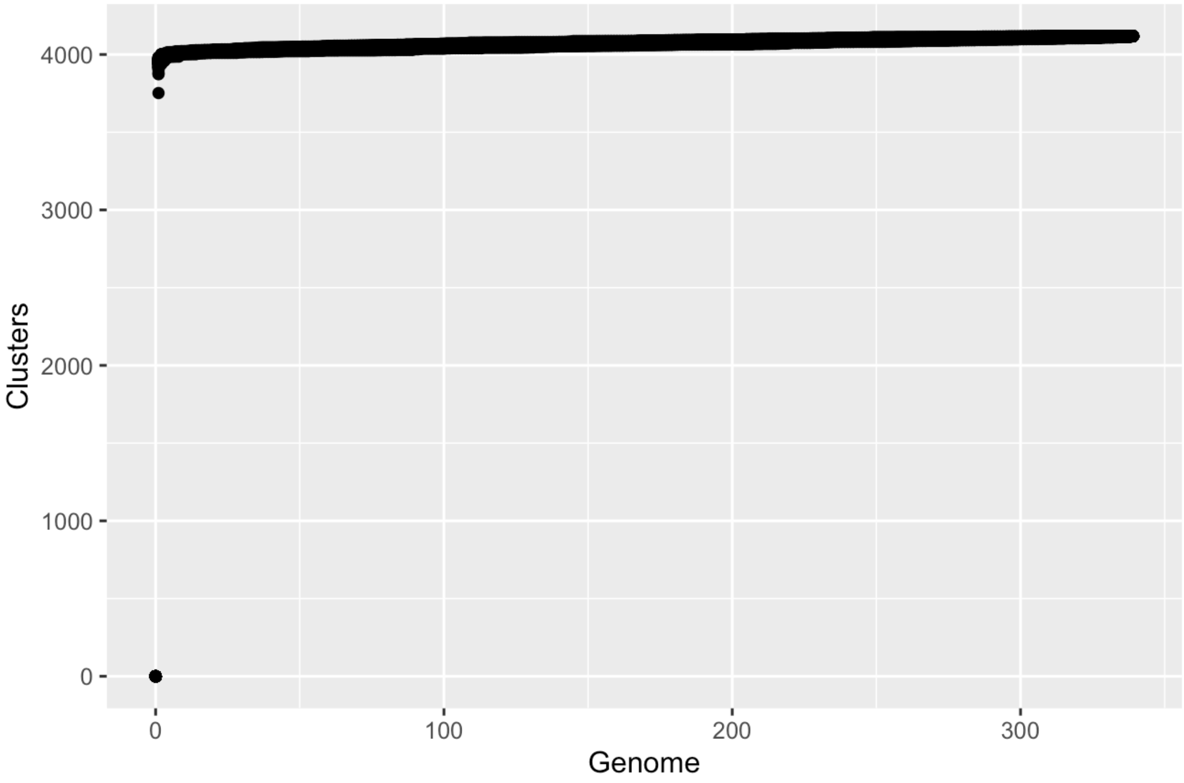

**B) Unmerged**:

| Fit to Heap's law: | Intercept | alpha |
|---|---|---|
| | 105.10826 | 1.548288 |

| Genome Fluidity: | Mean | Std |
|---|---|---|
| | 0.008819889 | 0.004459309 |

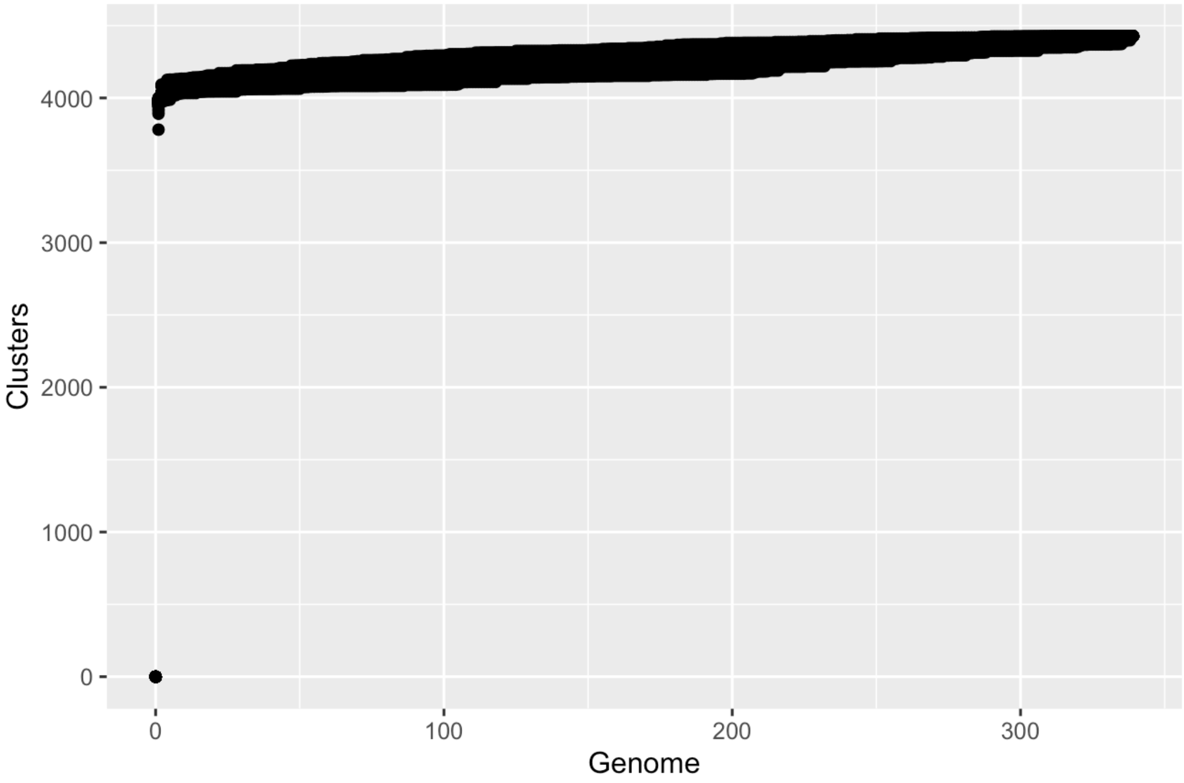

**C) Pangraph**:

| | | |
|---|---|---|
| **Fit to Heap's law:** | **Intercept** | **alpha** |
| | 89.29827 | 2 |
| **Genome Fluidity:** | **Mean** | **Std** |
| | 0.04136684 | 0.02142143 |

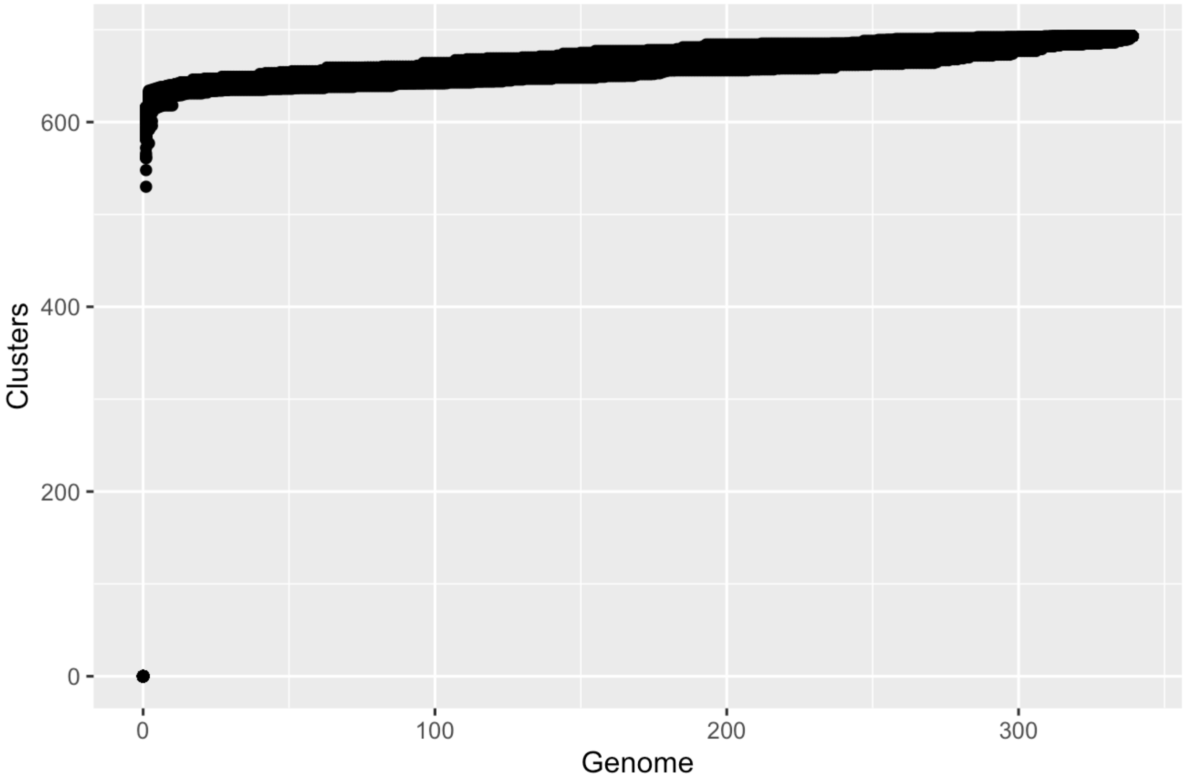
